# Membrane adaptation in the hyperthermophilic archaeon *Pyrococcus furiosus* relies upon a novel strategy involving glycerol monoalkyl glycerol tetraether lipids

**DOI:** 10.1101/2021.11.17.468962

**Authors:** Maxime Tourte, Philippe Schaeffer, Vincent Grossi, Philippe M. Oger

**Affiliations:** Univ Lyon, Univ. Lyon 1, CNRS, UMR 5240, F-69622, Villeurbanne, France; Univ Lyon, INSA Lyon, CNRS, UMR 5240, F-69621, Villeurbanne, France; Univ. Strasbourg, CNRS, UMR 7177, F-67000 Strasbourg, France; Univ Lyon, Univ. Lyon 1, CNRS, ENSL, UJM, UMR 5276 LGL-TPE, F-69622 Villeurbanne, France

**Keywords:** Archaeal membrane lipids, extremophiles, *Pyrococcus*, stress response, tetraethers

## Abstract

Microbes preserve membrane functionality under fluctuating environmental conditions by modulating their membrane lipid composition. Although several studies have documented membrane adaptations in Archaea, the influence of most biotic and abiotic factors on archaeal lipid compositions remains underexplored. Here, we studied the influence of temperature, pH, salinity, the presence/absence of elemental sulfur, the carbon source, and the genetic background on the core lipid composition of the hyperthermophilic neutrophilic marine archaeon *Pyrococcus furiosus.* Every growth parameter tested affected the core lipid composition to some extent, the carbon source and the genetic background having the greatest influence. Surprisingly, *P. furiosus* appeared to only marginally rely on the two major responses implemented by Archaea, i.e., the regulation of the ratio of diether to tetraether lipids and that of the number of cyclopentane rings in tetraethers. Instead, this species increased the ratio of glycerol monoalkyl glycerol tetraethers (GMGT, aka. H-shaped tetraethers) to glycerol dialkyl glycerol tetrathers (GDGT) in response to decreasing temperature and pH and increasing salinity, thus providing for the first time evidence of adaptive functions for GMGT. Besides *P. furiosus*, numerous other species synthesize significant proportions of GMGT, which suggests that this unprecedented adaptive strategy might be common in Archaea.

**Significance statement:** We describe here the membrane adaptive strategies the hyperthermophilic, neutrophilic, and marine model archaeon *Pyrococcus furiosus* implements in response to one of the largest sets of environmental stressors tested to date, including temperature, pH, salinity, presence/absence of elemental sulfur, carbon source, and genetic background. In contrast to the other archaea investigated so far, which response mainly involves the modulation of their diether/tetraether ratio and/or of their average number of cyclopentane rings, *P. furiosus* regulates its monoalkyl (so called H-shaped) to dialkyl tetraether ratio. Our study thus provides for the first time evidence of adaptive functions of archaeal monoalkyl tetraethers towards low temperature and pH and high salinity.

## Introduction

Membranes are essential compartments that control the cell inward and outward fluxes and support bioenergetic processes. These functions are however impeded by environmental conditions that microbes have to face in nature. To ensure proper membrane physicochemical properties and thus preserve cellular integrity and functions under contrasting conditions, Bacteria, Eukarya, and Archaea modulate their lipid compositions (Ernst *et al.*, 2016).

One of the most diagnostic features of Archaea compared to Bacteria and Eukarya is the unique structure of their membrane lipids. Indeed, instead of the lipids built upon straight fatty-acyl chains ester-bound to a glycerol backbone in a *sn*-1,2 configuration commonly found in Bacteria and Eukarya, Archaea synthesize lipids with polyisoprenoid alkyl chains that are ether-bound to a glycerol in a *sn*-2,3 configuration (De Rosa and Gambacorta, 1988). In addition to these typical bilayer-forming lipids, hereafter referred to as diether lipids, most Archaea are also capable of synthesizing membrane-spanning lipids, hereafter referred to as tetraether lipids, that generate monolayer membranes (De Rosa and Gambacorta, 1988). Recent technological advances in lipidomics further enlarged the diversity of lipid structures Archaea are able to synthesize, which now includes mono- and dialkyl glycerol diethers (MGD and DGD) with C20 and/or C25 alkyl chains (De Rosa *et al.*, 1986), glycerol mono-, di-, and trialkyl glycerol tetraethers (GMGT aka. H-GDGT, GDGT, and GTGT, respectively; (Morii *et al.*, 1998; Knappy *et al.*, 2011)), di- and tetraether lipids with hydroxylated and/or unsaturated isoprenoid chains (Gambacorta *et al.*, 1993; Nichols *et al.*, 2004), and tetraether lipids with glycerol, butanetriol, pentanetriol, and nonitol backbones (Becker *et al.*, 2016) (Figure 1). Besides this variety of lipid skeletons, or core lipids, Archaea also synthesize diverse polar head groups which mostly resemble those of Bacteria and Eukarya, i.e., phospho- and glyco-lipids deriving from sugars (glycerol, inositol, glucose, *N*-acetylhexosamine) (Jensen *et al.*, 2015), aminoacids (serine, ethanolamine) (Koga *et al.*, 1993), or combinations of both (Koga *et al.*, 1993) (Figure 1).

**Figure 1:**
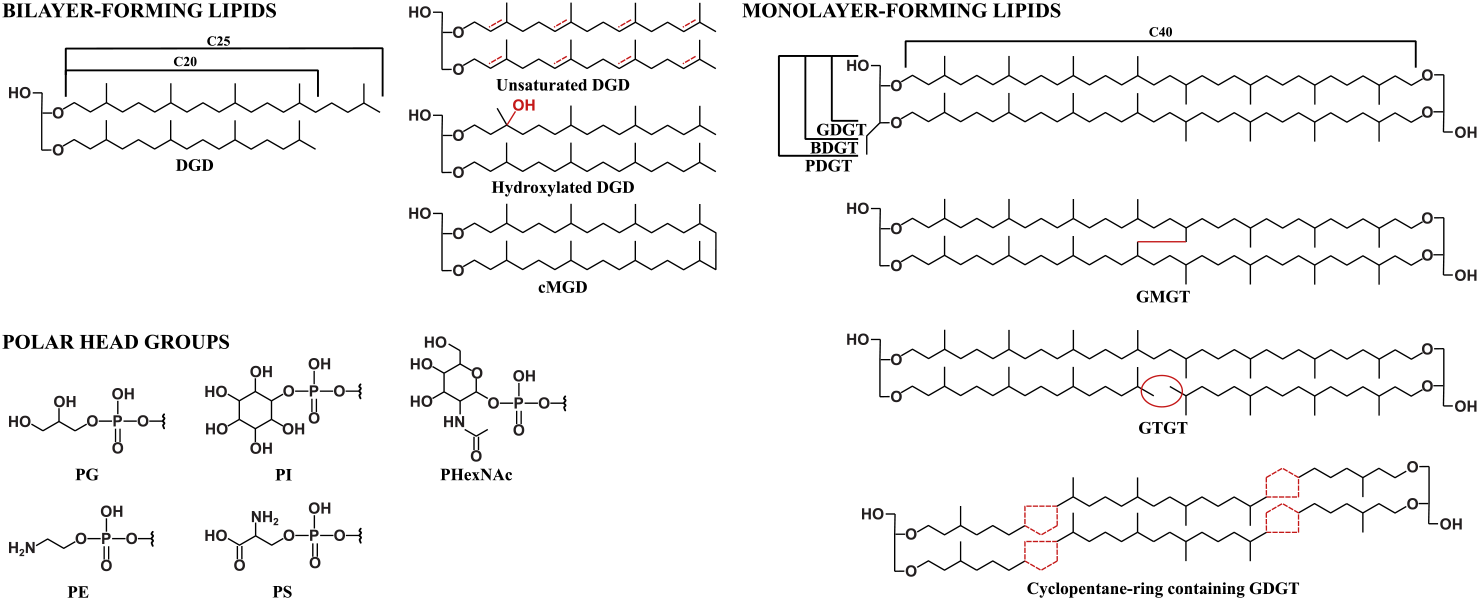
Archaeal membrane lipid diversity. Core archaeal lipids include dialkyl glycerol diethers (DGD) with C20 and/or C25 isoprenoid alkyl chains, unsaturated, and hydroxylated DGD, macrocyclic monoalkyl glycerol diethers (cMGD), glycerol, butanetriol and pentanetriol dialkyl glycerol tetraethers (GDGT, BDGT, and PDGT), glycerol monoalkyl glycerol tetraethers (GMGT), glycerol trialkyl glycerol tetraethers (GTGT), and tetraethers with 1 to 4 cyclopentane rings. Intact polar lipids consist of di- and tetraether core lipids attached to polar head groups deriving from sugars, e.g., phosphatidylinositol (PI), phosphatilyglycerol (PG), phosphatidyl-N-acetylhexosamine (PHexNAc), aminoacids, e.g., phosphoethanolamine (PE), and phosphatidylserine (PS), or combinations of both.

Archaea are the main inhabitants of the most severe environments on Earth, would it be due to temperature, pH, salinity, or hydrostatic pressure (Schleper *et al.*, 1995; Takai *et al.*, 2008; Birrien *et al.*, 2011), and this tolerance to multiple extreme conditions has been associated with the peculiar structure of their lipids. Indeed, polyisoprenoid alkyl chains provide greater membrane packing and impermeability compared to fatty acyl chains (Komatsu and Chong, 1998), while ether bonds are chemically and thermally more resistant than ester linkages (Baba *et al.*, 1999). Additionally, monolayer membranes generated by membrane-spanning tetraether lipids further enhance rigidity and impermeability compared to archaeal bilayers (Chong, 2010). While these physicochemical properties of classic diether and tetraether lipids rationalize the tolerance of Archaea to extreme conditions, the adaptive function and the behavior of membranes built upon the variety of lipids they can synthesize remain largely uncharted. Several membrane responses to abiotic stresses have nonetheless been evidenced in Archaea. The major membrane adaptation strategy in species producing a mixture of both di- and tetraether lipids consists in modulating the diether/tetraether ratio with changing growth conditions, which is congruent with the observation that monolayer-forming tetraether lipids tend to increase membrane packing, and thus stability, impermeability and rigidity compared to bilayer-forming diether lipids (Chong, 2010). In a pioneer work, Macalady and colleagues indeed reported increased proportions of tetraethers in archaeal species with lower pH optima (Macalady *et al.*, 2004). While such adjustments of the lipid compositions to optimal growth conditions do relate to adaptation, they do on a long time scale and are often not congruent with short-term adaptations – i.e., modifications of the lipid compositions in response to varying growth conditions – that were considered here. Decreased diether/tetraether ratios were observed in response to increasing temperature, decreasing hydrostatic pressure, or decreasing pH in a variety of Archaea (Sprott *et al.*, 1991; Lai *et al.*, 2008; Matsuno *et al.*, 2009; Cario *et al.*, 2015; Jensen *et al.*, 2015). In contrast, the membrane response of species producing only tetraether lipids involves the fine regulation of the number of unsaturations – in the form of cyclopentane rings – present along the hydrophobic alkyl chains in response to temperature and pH (Shimada *et al.*, 2008; Elling *et al.*, 2014; Jensen *et al.*, 2015; Bale *et al.*, 2019), according to the rationale that a higher number of rings induces a more compact membrane and, hence, enhances stability, impermeability and rigidity (Gliozzi *et al.*, 1983; Gabriel and Chong, 2000). However, whereas Archaea living at lower pH do tend to have higher numbers of rings (long term), there are examples of strains decreasing their ring index in response to decreased pH (short term; see for instance *Thermoplasma acidophilum* in (Shimada *et al.*, 2008)). In a similar manner, the membrane homeostasis of species at the opposite side of the lipid spectrum, i.e., producing exclusively diether lipids, also involves the regulation of the lipid unsaturation levels – here in the form of double bonds (Nichols *et al.*, 2004; Gibson *et al.*, 2005; Dawson *et al.*, 2012). For instance, the psychrophilic archaeon *Methanococcoides burtonii* increases its relative proportions of unsaturated diether lipids in response to decreased temperature (Nichols *et al.*, 2004), in a way reminiscent of the typical adaptive strategy employed by Eukaryotes and Bacteria (Grossi *et al.*, 2010; Ernst *et al.*, 2016). Last, in addition to typical diether and tetraether lipids, the hyperthermophilic methanogen *Methanocaldococcus jannaschii* synthesizes an intriguing macrocyclic diether lipid in which the two phytanyl chains are covalently linked *via* a C-C bond, hereafter referred to as cMGD (Figure 1; (Comita *et al.*, 1984)). This covalent bond reduces the lateral motion of the cMGD and increases membrane packing, stability and impermeability to solutes and protons (Dannenmuller *et al.*, 2000; Arakawa *et al.*, 2001). The relative proportions of cMGD in *M. jannaschii* was thus naturally observed to increase in response to increased temperature (Sprott *et al.*, 1991). While all the aforementioned membrane responses concerned typical abiotic factors, Archaea were also demonstrated to modulate their lipid compositions in response to numerous (if not all) biotic parameters, e.g., carbon, phosphorous, and nitrogen sources and availability (Langworthy, 1977; Matsuno *et al.*, 2009; Elling *et al.*, 2014; Meador *et al.*, 2014; Feyhl-Buska *et al.*, 2016; Quehenberger *et al.*, 2020; Zhou *et al.*, 2020), although such reports remain scarce. The relative contribution of the polar moieties of the ether lipids in the membrane adaptation of Archaea remains in contrast underexplored.

*Pyrococcus furiosus* is a model archaeon belonging to the Thermococcales order that has been isolated from geothermally heated sediments on the coast of Vulcano Island, Italy (Fiala and Stetter, 1986), which are naturally subjected to contrasting conditions due to changes in tide and geothermal regimes (Rogers *et al.*, 2007). The growth of *P. furiosus* occurs by fermentation of peptide and/or sugar mixtures under a large range of growth conditions, including temperature from 70 to 103 °C (optimal 98 °C), pH from 5 to 9 (optimal 6.8), and salinity from 0.5 to 5 % *w/v* of NaCl (optimal 3 % NaCl), which allows the description of its adaptive strategy towards a large spectrum of biotic and abiotic parameters. In addition, preliminary investigations of *P. furiosus* membrane content revealed a unique lipid composition (Figure S1). While it exhibits a limited diversity of polar head groups, i.e., phosphoinositol and a few derivatives (Sprott *et al.*, 1997; Lobasso *et al.*, 2012; Tourte *et al.*, 2020a), those are attached to as many as 14 different core structures, namely DGD, GDGT with 0 to 4 cyclopentane rings (GDGT0-4, respectively), GTGT with 0 to 2 cyclopentane rings (GTGT0-2, respectively), and GMGT with 0 to 4 cyclopentane rings (GMGT0-4) (Tourte *et al.*, 2020a) (Figure S1), paving the way for the elucidation of uncommon membrane adaptive strategies. Indeed, we show here that *P. furiosus* modulates the ratio of GMGT/GDGT and marginally regulates the number of cyclopentane rings instead of altering the diether/tetraether ratio as most other Archaea do, to respond to temperature, pH, salinity, the presence/absence of elemental sulfur, growth medium, and genetic background. In particular, the relative proportions of GMGT were significantly increased at low pH and high salinity, which highlights for the first time the adaptive functions of these intriguing core lipid structures to pH and salinity in a hyperthermophilic archaeon.

## Results

### *Pyrococcus furiosus* displays a large core lipid diversity under contrasting growth conditions

Although the hyperthermophilic archaeon *Pyrococcus furiosus* DSM3638 is able to grow over a wide range of conditions (Fiala and Stetter, 1986), only some of them allow growth yields (> 10^8^ cell ml^-1^) and rates (> 0.08 h^-1^) compatible with lipid analysis. For instance, temperatures below 80 °C were not assessed despite *P. furiosus* being theoretically able to grow down to 70 °C. As numerous biotic parameters have been shown to impact lipid compositions (e.g., growth stage (Elling *et al.*, 2014)), growth was tightly monitored (refer to Table S1 for growth measurements under the conditions tested here) and cultures were all inoculated at a cell density of 10^5^ cell ml^-1^ and harvested in late log phase so as to limit the influence of other parameters on the lipid compositions observed.

*P. furiosus* DSM3638 was previously shown to produce at least 14 different core lipid structures (Figure S1) under optimal growth conditions, i.e., in Thermococcales rich medium (TRM) at 98 °C and pH 6.8, with 3 % *w/v* NaCl and 10 g L^-1^ of elemental sulfur (Tourte *et al.*, 2020a). Nine core structures, i.e., DGD, GDGT with up to 3 cyclopentane rings (GDGT0 to 3), GTGT without cyclopentane ring (GTGT0), and GMGT with up to 3 cyclopentane rings (GMGT0 to 3) were detected in all conditions tested and constituted the core set of core lipid structures, whereas the five remaining core lipids, i.e., GDGT with 4 cyclopentane rings (GDGT4; up to 3.5 %), GTGT with 1 or 2 cyclopentane rings (GTGT1 and 2; up to 2.4 and 1.0 %, respectively), and GMGT with 4 cyclopentane rings (GMGT4; up to 0.8 %), were only detected in low proportions under specific conditions (Table 1), and were thus considered as accessory or minor lipids.

**Table 1.**
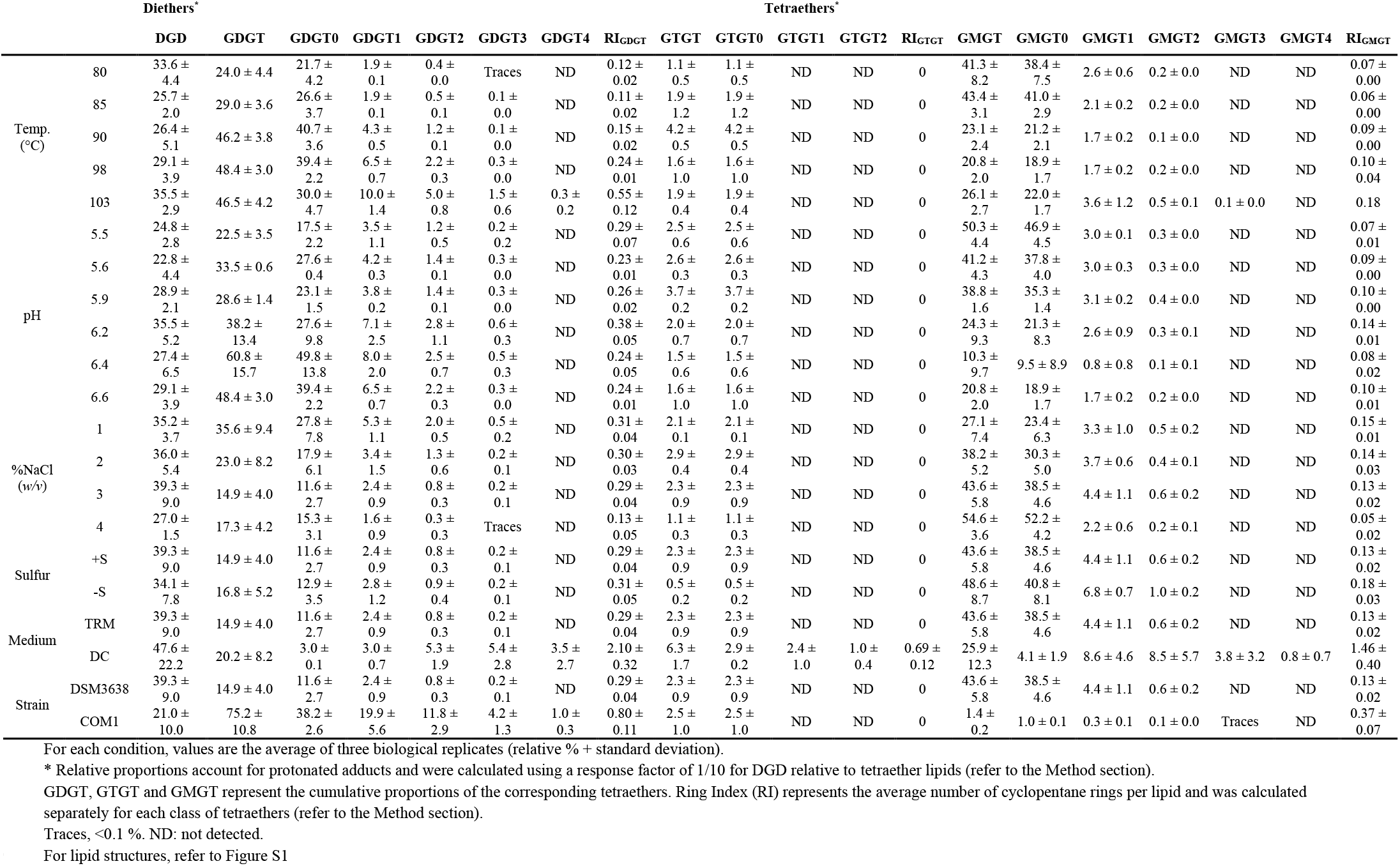
Core lipid relative composition (%) of *Pyrococcus furiosus* as a function of growth variables.

### Impact of temperature on the core lipid composition of *P. furiosus*

The effect of growth temperature on the membrane core lipid composition of *P. furiosus* was investigated with cultures grown at 80, 85, 90, 98 (T_opt_), and 103 °C. The same core lipid structures, i.e., DGD, GDGT0 to 3, GTGT0, and GMGT0 to 3 were detected under all temperatures tested, with the exception of GDGT4 which was detected only when *P. furiosus* was grown at 103 °C (Table 1, Figure 2A). To better account for the diversity of GDGT and GMGT structures, the proportions of all derivatives from each class of tetraether lipids (GDGT, GMGT) were summed and their diversity was represented by their average number of rings per molecule, or ring index (RI). Different RI were calculated to evaluate the incorporation of cyclopentane rings in the different tetraether populations, namely RITetraethers which represents the average number of cyclopentane rings in all tetraethers, and RIGDGT and RIGMGT which represent the average number of cyclopentane rings in GDGT and GMGT, respectively (see methods for the calculation formulas).

**Figure 2:**
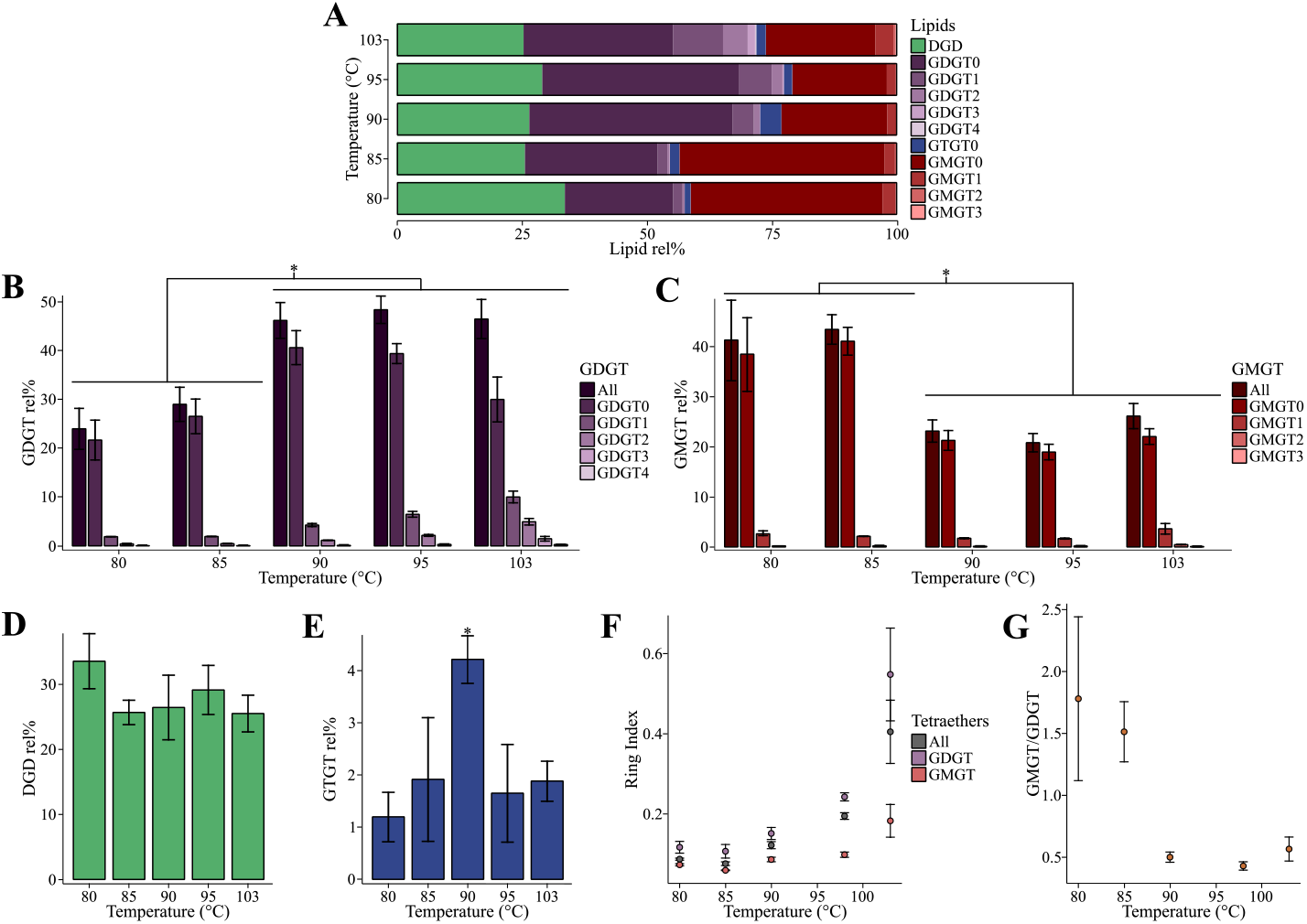
*Pyrococcus furiosus* responds to increasing temperature by increasing the average number of cyclopentane rings. *P. furiosus* DSM3638 was grown under optimal conditions (TRM at 3 % *w/v* NaCl and pH 6.8 with 10 g L^-1^ elemental sulfur) at different temperatures (80, 85, 90, 98, and 103 °C). 98 °C represents the optimal growth temperature. Error bars represent the standard deviation calculated on three biological replicates. * indicates temperatures with significantly different core lipid compositions. (A) Total core lipid compositions under each temperature. (B) Influence of the temperature on GDGT relative proportions. GDGT (dark purple) corresponds to the summed GDGT, regardless of the cyclopentane ring content. (C) Influence of the temperature on GMGT relative proportions. GMGT (dark red) corresponds to the summed GMGT, regardless of the cyclopentane ring content. (D) Influence of the temperature on DGD relative abundance. (E) Influence of the temperature on GTGT0 relative abundance. (F) Temperature dependence of the ring index (RI) for all tetraethers (GDGT, GTGT, and GMGT), GDGT and GMGT (RI ± standard deviation). (G) Temperature dependence of the GMGT/GDGT ratio.

Small differences were observed in the DGD and GTGT contents under the distinct growth temperatures tested. The relative proportions of DGD ranged from 25.7 ± 2.0 % at 85 °C to 33.6 ± 4.4 % at 80 °C and those of GTGT0 from 1.2 ± 0.4 % at 80 °C to 4.2 ± 0.5 % at 90 °C (Table 1, Figures 2D and 2E). In contrast, the GDGT and GMGT contents showed much larger variations. The two lowest temperatures (80 and 85 °C) and the three highest (90, 98 and 103 °C) differed significantly from one another based on their GDGT (~25-30 % *vs*. ~50 %) and GMGT (~40 % *vs*. ~20-25 %) compositions (Table 1, Figures 2B and 2C), which would suggest the existence of a threshold temperature above which GMGT are substituted by GDGT. This is best illustrated when representing the ratio between GMGT and GDGT (GMGT/GDGT) which dropped from 1.78 ± 0.67 and 1.51 ± 0.25 at 80 and 85°C to 0.50 ± 0.5, 0.43 ± 0.04 and 0.57 ± 0.10 at 90, 98, and 103 °C, respectively (Table 1, Figure 2G). The membrane response of *P. furiosus* DSM3638 to temperature is further completed by variations of the RIGMGT and RIGDGT. Although all the different RI increased with temperature, they showed contrasting trends. Whereas RIGDGT and RITetraethers continuously increased from 0.11 ± 0.02 and 0.08 ± 0.01 at 85 °C to 0.55 ± 0.12 and 0.40 ± 0.08 at 103 °C, respectively (Table 1, Figure 2F), RI_GMGT_ values were constant at *ca.* 0.06-0.10 ± 0.01 from 85 °C to 98°C, and only increased significantly at 103 °C to reach 0.18 ± 0.04.

Core lipid compositions and temperatures were correlated using the Spearman’s coefficient ρ (Table 2). Although a clear critical threshold can be drawn between 85 and 90 °C, no clear trend was observed between the GMGT/GDGT ratio and temperature. No significant correlation between DGD, GTGT, and GMGT proportions and temperature was observed, but GDGT, and especially cyclopentane ring-containing GDGT, were significantly positively correlated with temperatures (ρ = +0.90, +1.00, +1.00 and +1.00 for all GDGT, GDGT1, GDGT2 and GDGT3, respectively). As a consequence, RI_GDGT_ (ρ = +0.90) and RI_Tetraethers_ (ρ = +0.90) were also significantly positively correlated with temperature. This was also valid for RI_GMGT_ (ρ = +0.90), although no clear trend could be highlighted between temperature and the proportion of GMGT with and without cyclopentane rings.

**Table 2.** Spearman correlation coefficient (ρ) and P-values for individual core structures and total GDGT, GTGT and GMGT as a function of temperature, pH and salinity.

|  | Temperature (°C) |  | pH |  | %NaCl (%w/v) |  |
| --- | --- | --- | --- | --- | --- | --- |
| | Spearman's $\rho$ | P-value | Spearman's $\rho$ | P-value | Spearman's $\rho$ | P-value |
| DGD | -0.60 |  | 0.66 |  | -0.2 |  |
| GDGT | 0.90 | * | 0.89 | * | -0.8 |  |
| GDGT0 | 0.60 |  | 0.89 | * | -0.8 |  |
| GDGT1 | 1.00 | **** | 0.77 |  | -1 | **** |
| GDGT2 | 1.00 | **** | 0.77 |  | -1 | **** |
| GDGT3 | 1.00 | **** | 0.77 |  | -1 | **** |
| GDGT4 | 0.71 |  | NA | NA | NA | NA |
| GTGT | 0.2 |  | -0.71 |  | -0.4 |  |
| GTGT0 | 0.2 |  | -0.71 |  | -0.4 |  |
| GTGT1 | NA | NA | NA | NA | NA | NA |
| GTGT2 | NA | NA | NA | NA | NA | NA |
| GMGT | -0.60 |  | -0.94 | ** | 1 | **** |
| GMGT0 | -0.60 |  | -0.94 | ** | 1 | **** |
| GMGT1 | 0 |  | -0.71 |  | -0.2 |  |
| GMGT2 | 0.20 |  | -0.54 |  | -0.4 |  |
| GMGT3 | 0.71 |  | NA |  | NA | NA |
| GMGT4 | NA | NA | NA |  | NA | NA |

NA, not applicable because the core structure was not detected.
Only P-values $\leq 0.05$ are indicated, as follows: < 0.0001, "\*\*\*\*"; 0.0001 to 0.001, "\*\*\*\*", 0.001 to 0.01, "\*\*\*" and 0.01 to 0.05, "\*\*".

### Effect of pH on the core lipid composition of *P. furiosus*

Although *Pyrococcus furiosus* DSM3638 has been reported to be able to grow from pH 5.0 to 9.0, we could only assess its membrane adaptation to mild acidic pH (from 5.0 to 6.8) due to the instability of alkaline buffers at temperatures close to 100 °C. Under all the tested pH, the same set of core lipid structures, i.e., DGD, GDGT0 to 3, GTGT0, and GMGT0 to 3 was found (Table 1, Figure 3A). As seen for the temperature experiments, small differences in DGD and GTGT0 contents were observed, whereas the proportions of GDGT and GMGT showed much larger variations. Indeed, the relative proportions of DGD ranged from 22.8 ± 4.4 % at pH 5.6 to 35.6 ± 5.2 % at pH 6.2 and those of GTGT0 varied from 1.5 ± 0.6 % at pH 6.4 to 3.7 ± 0.2 % at pH 5.9 (Table 1, Figures 3D and 3E), whereas GDGT and GMGT showed relative abundances ranging from 22.5 ± 3.5 % and 50.3 ± 4.4 % at pH 5.5 to 60.8 ± 15.7 % to 10.3 ± 9.7 % at pH 6.4, respectively (Table 1, Figures 3B and 3C). In a manner similar to what was observed for temperature, two distinct groups could be delineated: cultures grown at pH 5.5, 5.6, and 5.9 had more GMGT than GDGT, and thus GMGT/GDGT ratios above 1 (i.e., 2.28 ± 0.47, 1.23 ± 0.14, and 1.36 ± 0.10, respectively), whereas cultures grown at higher pH, i.e. 6.2, 6.4, and 6.6, had more GDGT than GMGT and displayed GMGT/GDGT ratios below 1 (i.e., 0.77 ± 0.57, 0.21 ± 0.25 and 0.43 ± 0.04, respectively; Table 1, Figure 3G). In contrast to the response to temperature though, no clear trend could be observed in the RI values regardless of the class of tetraether (Table 1, Figure 3F). Altogether, these results suggest that pH mostly alters the GMGT/GDGT ratio in *P. furiosus.* This was further supported by the Spearman’s correlations (Table 2): while the summed GDGT and GDGT0 relative proportions significantly increased with pH (ρ = +0.89 for both), those of the summed GMGT and GMGT0 showed an opposite trend (ρ = −0.94 for both), which resulted in a GMGT/GDGT ratio significantly negatively correlated with pH (ρ = −0.89). In contrast, none of the DGD, GTGT, or ring-containing structures, and thus none of the RI, significantly correlated with pH.

**Figure 3:**
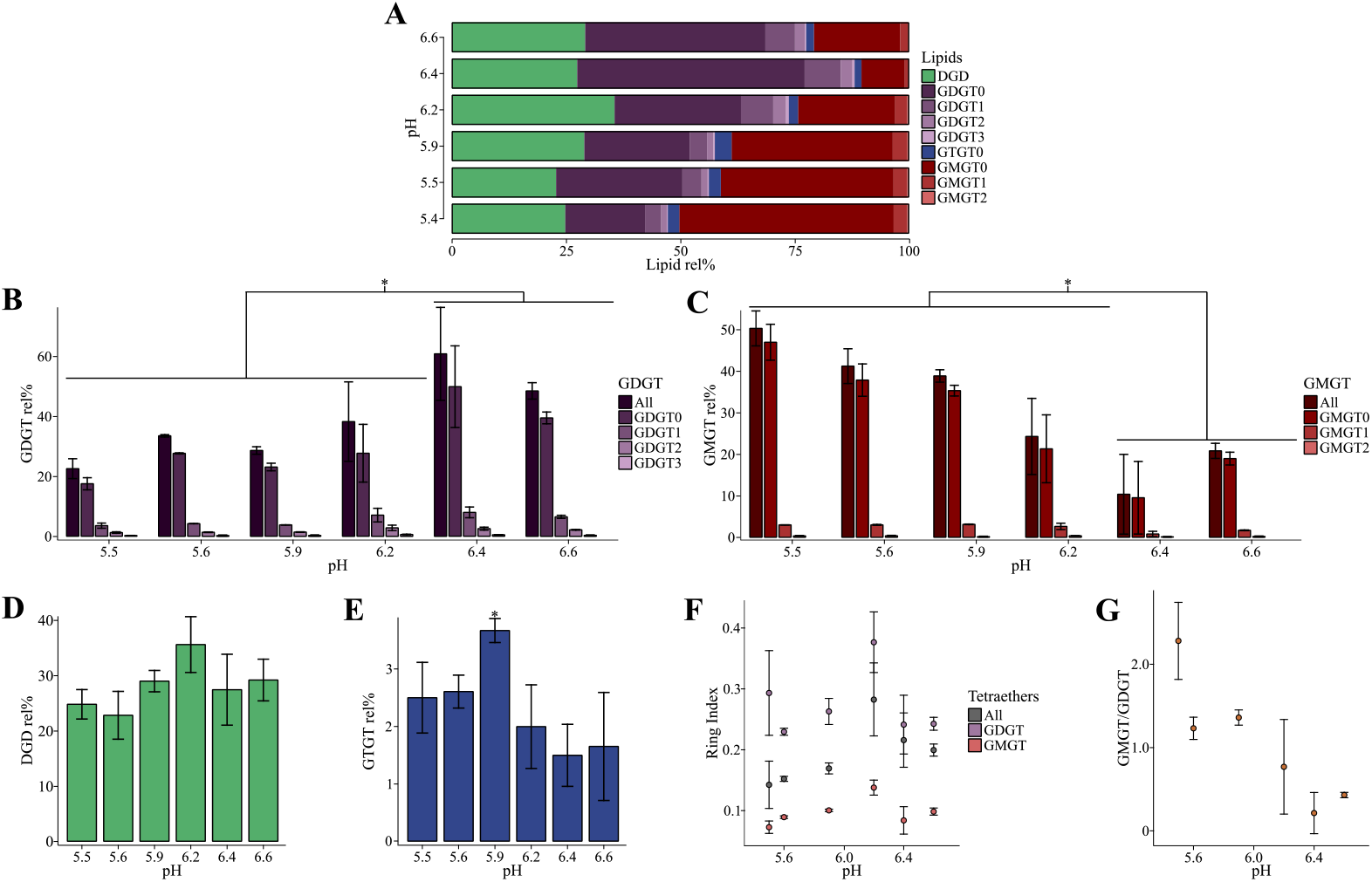
*Pyrococcus furiosus* responds to sub-optimal pH by increasing the relative abundance of GMGT. *P. furiosus* DSM3638 was grown under optimal conditions (TRM at 3 % *w/v* NaCl and 98 °C with 10 g L^-1^ elemental sulfur) at different pH (5.5, 5.6, 5.9, 6.2, 6.4 and 6.6). pH 6.6 represents the optimal growth pH. Error bars represent the standard deviation calculated on three biological replicates. * indicates pH with significantly different core lipid compositions. (A) Total core lipid compositions under each pH. (B) Influence of the pH on GDGT relative proportions. GDGT (dark purple) corresponds to the summed GDGT, regardless of the cyclopentane ring content. (C) Influence of the pH on GMGT relative proportions. GMGT (dark red) corresponds to the summed GMGT, regardless of the cyclopentane ring content. (D) Influence of the pH on DGD relative abundance. (E) Influence of the pH on GTGT0 relative abundance. (F) pH dependence of the ring index (RI) for all tetraethers (GDGT, GTGT, and GMGT), GDGT and GMGT (RI ± standard deviation). (G) pH dependence of the GMGT/GDGT ratio.

### Effect of salinity on the core lipid composition of *P. furiosus*

We tested the membrane response of *P. furiosus* strain DSM3638 to salinity covering from 1 to 4 % *w/v* of NaCl, as salinities outside this range resulted in extremely low growth yields. Cultures grown under all salinities tested exhibited the same set of core lipids, i.e., DGD, GDGT0 to 3, GTGT0 and GMGT0 to 2 (Table 1, Figure 4A). As for temperature and pH, small differences in the DGD and GTGT0 contents were observed, the relative proportions of DGD ranging from 27.0 ± 1.5 % at 4 % NaCl to 39.3 ± 9.0 % at 3 % NaCl and those of GTGT0 from 1.1 ± 0.3 % at 4 % NaCl to 2.9 ± 0.4 % at 2 % NaCl (Table 1, Figures 4D and 4E). Only the GTGT0 content at 4 % NaCl appeared significantly distinctive *(ca.* 1.1 % *vs.* above 2.0 %). Similarly to the aforementioned stresses, the relative abundances of GDGT and GMGT were more affected by changes in salinity and ranged from 35.6 ± 9.4 % and 27.1 ± 7.4 % at 1 % NaCl to 17.3 ± 4.2 % and 54.6 ± 3.6 % at 4 % NaCl, respectively (Table 1, Figures 4B and 4C). The GMGT/GDGT ratio thus increased with increasing salinity, with values ranging from 0.83 ± 0.37 at 1 % NaCl to 3.30 ± 0.92 at 4 % NaCl, respectively (Table 1, Figure 4G). In contrast to pH and temperature, increasing salinity tended to decrease the RI_GDGT_, RI_GMGT_ and RI_Tetraethers_, with values ranging from 0.31 ± 0.04, 0.15 ± 0.01, and 0.23 ± 0.02 at 1 % NaCl to 0.13 ± 0.04, 0.05 ± 0.02, and 0.07 ± 0.03 at 4 % NaCl, respectively (Table 1, Figure 4F). The summed GMGT and GMGT0 (ρ = +1.00 for both) and the GMGT/GDGT ratio (ρ = +1.00) were significantly positively correlated with the % NaCl, whereas significant negative correlations were observed between the % NaCl and GDGT1 to 3 and RI_GDGT_, RI_GMGT_ and RI_Tetraethers_ (ρ = −1.00 for all) (Table 2). Thus, the major alterations triggered by salinity on the core lipid composition of *P. furiosus* appeared to be the fine tuning of the GMGT/GDGT ratio and of the number of cyclopentane rings.

**Figure 4:**
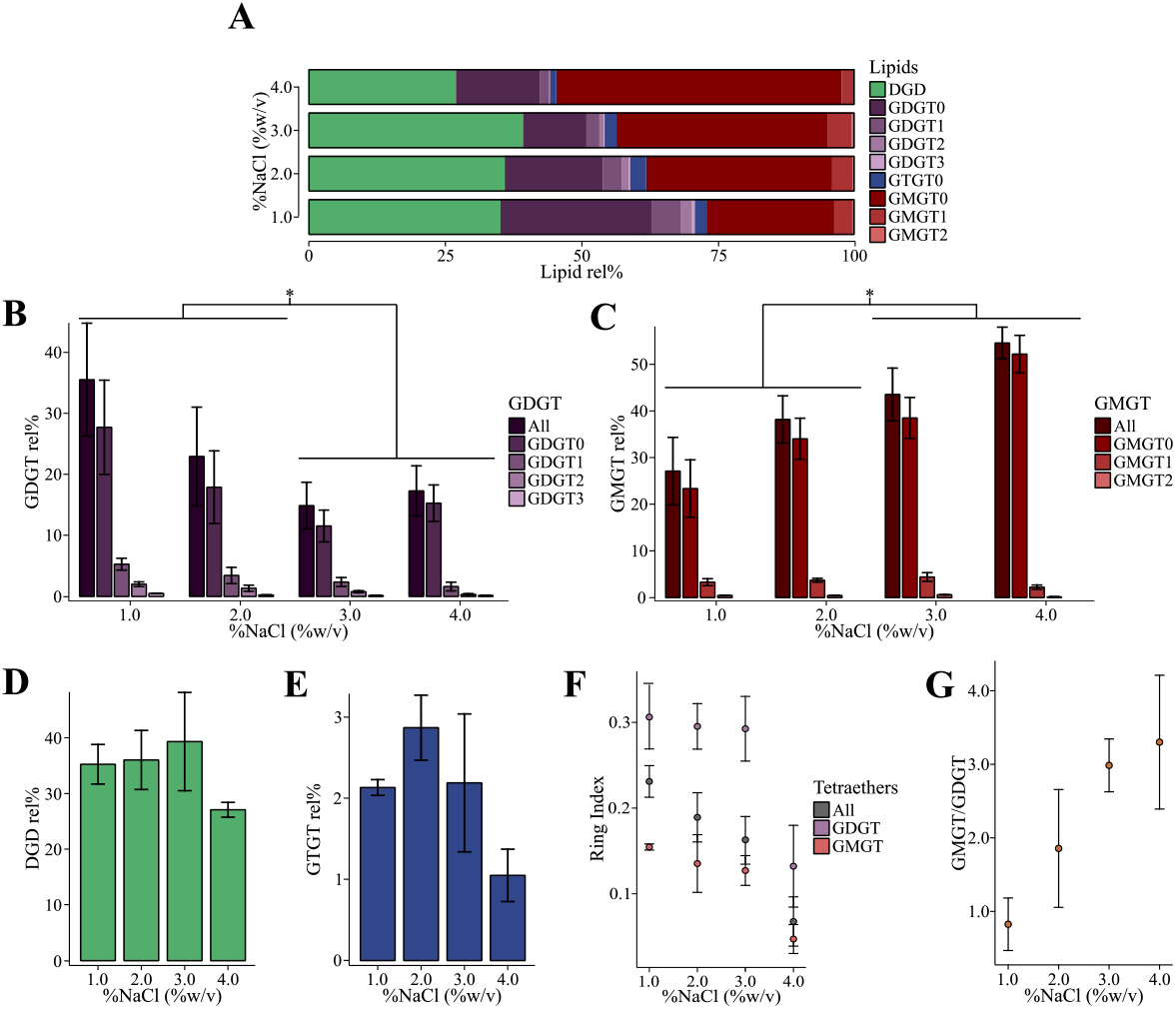
*Pyrococcus furiosus* responds to increasing salinity by increasing the relative abundance of GMGT and reducing the average number of cyclopentane rings. *P. furiosus* DSM3638 was grown under optimal conditions (TRM at 98 °C and pH 6.8 with 10 g L^-1^ elemental sulfur) at different salinities (1, 2, 3 and 4 % *w/v* NaCl). 3 % NaCl represents the optimal growth salinity. Error bars represent the standard deviation calculated on three biological replicates. * indicates % NaCl with significantly different core lipid compositions. (A) Total core lipid compositions under each % NaCl. (B) Influence of the salinity on GDGT relative proportions. GDGT (dark purple) corresponds to the summed GDGT, regardless of the cyclopentane ring content. (C) Influence of the salinity on GMGT relative proportions. GMGT (dark red) corresponds to the summed GMGT, regardless of the cyclopentane ring content. (D) Influence of the salinity on DGD relative abundance. (E) Influence of the salinity on GTGT0 relative abundance. (F) Salinity dependence of the ring index (RI) for all tetraethers (GDGT, GTGT, and GMGT), GDGT and GMGT (RI ± standard deviation). (G) Salinity dependence of the GMGT/GDGT ratio.

### Impact of the presence/absence of elemental sulfur on the core lipid composition of *P. furiosus*

Like other Thermococcales, *P. furiosus* uses sulfur to detoxify H2, a major by-product of its heterotrophic metabolism (Chou *et al.*, 2007). Although elevated concentrations of H2 in closed cultures are toxic for some Thermococcales, significant growth of *P. furiosus* can be achieved without the addition of sulfur (Fiala and Stetter, 1986). The absence of elemental sulfur nonetheless requires a lag time for adaptation, which suggests that the lack of elemental sulfur is perceived as a stress (Table S1). Here, we investigated the influence of the presence or absence of elemental sulfur on *P. furiosus* core lipid composition by assessing two concentrations: 0 (-S) and 10 g L^-1^ (saturation level; +S). Under such growth conditions, the same core lipid structures, i.e., DGD, GDGT0 to 3, GTGT0 and GMGT0 to 2, could be identified (Table 1, Figure S2A). In contrast to all the other parameters tested, the core lipid compositions were not significantly different in the presence or absence of elemental sulfur. For instance, growth with sulfur resulted in relative proportions of DGD, GDGT, and GMGT of 39.3 ± 9.0, 14.9 ± 4.0, and 43.6 ± 5.8 %, respectively, whereas those after growth without sulfur were of 34.1 ± 7.8, 16.8 ± 5.2, and 48.6 ± 8.7 %, respectively (Table 1, Figure S2B-D). Similarly, no significant differences were observed for the GMGT/GDGT ratio and the RI values (Table 1, Figure S2F-G). Only the relative proportions of GTGT0 were significantly different between the two conditions, i.e., 2.2 ± 0.9 with and 0.5 ± 0.1 % without elemental sulfur (Table 1, Figure S2E).

### Impact of the carbon source on the core lipid composition of *P. furiosus*

It is well established in Bacteria and Eukaryotes that the carbon source greatly impacts membrane lipid compositions (Vinçon-Laugier *et al.*, 2016). We tested the impact of the switch from a proteinaceous carbon source in TRM medium to a reducing sugar, namely cellobiose, in defined cellobiose (DC) medium on the lipid composition of *P. furiosus* DSM3638. Surprisingly, in addition to the core lipids typically synthesized by the strain, i.e., DGD, GDGT0 to 4, GTGT0, and GMGT0 to 3, novel core structures, i.e., GTGT1 and 2, and GMGT4, were identified during growth in DC medium (Table 1, Figure 5A). Despite these additional core lipid structures, the relative abundances of DGD (47.6 ± 22.2 *vs*. 39.3 ± 9.0), summed GDGT (14.9 ± 4.0 *vs*. 20.2 ± 8.2), and summed GMGT (43.6 ± 5.8 *vs*. 25.9 ± 12.3) were not significantly different between DC and TRM media, respectively (Table 1, Figure 5B-D). When grown in DC medium, *P. furiosus* nonetheless harbored significantly higher proportions of GDGT1 to 4 and of GMGT1 to 4, which resulted in much higher values for RI_GDGT_, RI_GMGT_, and RI_Tetraethers_, i.e., 2.10 ± 0.32, 1.46 ± 0.40, and 1.61 ± 0.34 in DC compared to 0.29 ± 0.04, 0.13 ± 0.02, and 0.16 ± 0.03 in TRM, respectively (Table 1, Figure 5F). In contrast to any other growth condition tested here, the nature of the growth medium impacted the total proportions of GTGT, which was significantly higher in DC (6.3 ± 1.7) than in TRM medium (2.3 ± 0.9), notably due to the presence of GTGT1 (2.4 ± 1.0) and GTGT2 (1.0 ± 0.4) that were not detected in TRM medium (Table 1, Figure 5E). Similarly to all the RI calculated previously, RIGTGT was thus much higher in DC than in TRM medium (0.69 ± 0.12 *vs.* 0; Table 1, Figure 5F). The major core lipid composition changes triggered by the switch in carbon source from protein to sugar thus appeared mostly restricted to cyclopentane ring-containing tetraethers.

**Figure 5:**
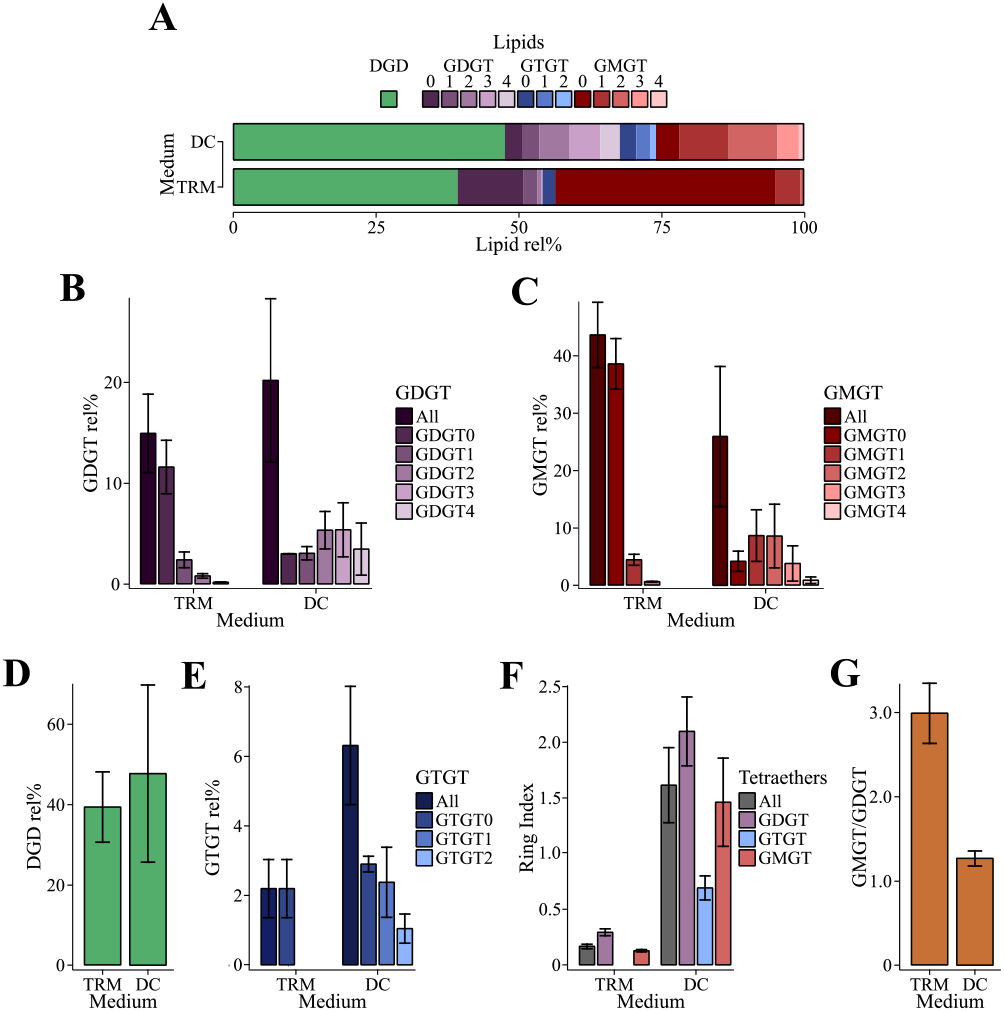
Growth on sugars significantly increases the average number of cyclopentane rings in *Pyrococcus furiosus*. *P. furiosus* DSM3638 was grown under optimal conditions (at 98 °C, pH 6.8 and 3 % *w/v* NaCl with 10 g L^1^ elemental sulfur) in DC and TRM media. TRM represents the optimal growth condition. For details on media compositions, refer to the Method section. Error bars represent the standard deviation calculated on three biological replicates. (A) Total core lipid compositions under each medium condition. (B) Influence of the growth medium on GDGT relative proportions. GDGT (dark purple) corresponds to the summed GDGT, regardless of the cyclopentane ring content. (C) Influence of the growth medium on GMGT relative proportions. GMGT (dark red) corresponds to the summed GMGT, regardless of the cyclopentane ring content. (D) Influence of the growth medium on DGD relative abundance. (E) Influence of the growth medium on GTGT0 relative abundance. (F) Growth medium dependence of the ring index (RI) for all tetraethers (GDGT, GTGT, and GMGT), GDGT, GTGT and GMGT (RI ± standard deviation). (G) Growth medium dependence of the GMGT/GDGT ratio.

### Lipid composition of the quasi-isogenic *P. furiosus* strain COM1

We tested whether small genetic modifications could induce variations in the lipid compositions of near isogenic strains of *P. furiosus,* i.e., the wild type strain DSM3638 and the genetically tractable COM1 strain which is a *pyrF*-deleted derivative of DSM3638 (Bridger *et al.*, 2012). As expected, the two isolates exhibited the same set of core lipids structures, i.e., DGD, GDGT0 to 3, GTGT0, and GMGT0 to 3, but in surprisingly different proportions (Table 1, Figure 6A). Indeed, strains COM1 and DSM3638 showed notably different GDGT (75.2 ± 10.8 % *vs*. 14.9 ± 4.0 %) and GMGT (1.4 ± 0.2 % *vs*. 43.6 ± 5.8) contents, which resulted in very contrasting GMGT/GDGT ratios (0.02 ± 0.01 *vs*. 2.99 ± 0.37; Table 1, Figures 6B, 6C and 6G). The higher total proportion of GDGT observed for strain COM1 reflects a significant increase in the abundance of all ring-containing GDGT structures, including GDGT4, a core structure that was not detected in strain DSM3638 under optimal growth conditions (Table 1, Figure 6B). Although strain COM1 exhibited minor proportions of GMGT, ring-containing structures also appeared in comparatively higher proportions. The RI_GDGT_, RI_GMGT_, and RI_Tetraethers_ were consequently higher for COM1, with values of 0.80 ± 0.11, 0.37 ± 0.07, and 0.76 ± 0.12 for strain COM1 compared to 0.29 ± 0.04, 0.13 ± 0.02, and 0.16 ± 0.03 for strain DSM3638, respectively (Table 1, Figure 6F).

**Figure 6:**
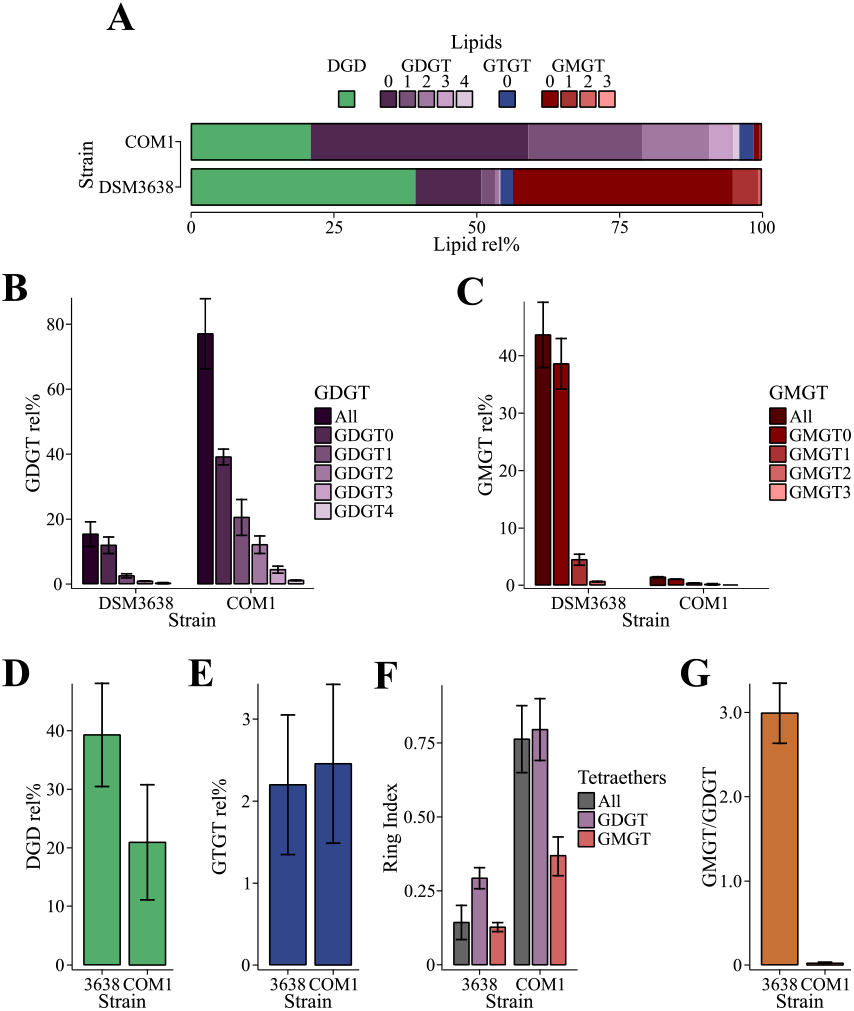
*Pyrococcus furiosus* strain COM1 shows a marked increase in GDGT content and average number of cyclopentane rings compared to its wild-type parent strain DSM3638. Cells were grown under optimal conditions (TRM at 98 °C, pH 6.8 and 3 % *w/v* NaCl with 10 g L^1^ elemental sulfur). Error bars represent the standard deviation calculated on three biological replicates. (A) Total core lipid compositions for each strain. (B) GDGT relative abundance for each strain. GDGT (dark purple) corresponds to the summed GDGT, regardless of the cyclopentane ring content. (C) GMGT relative abundance for each strain. GMGT (dark red) corresponds to the summed GMGT, regardless of the cyclopentane ring content. (D) DGD relative abundance for each strain. (E) GTGT relative abundance for each strain. (F) Strain dependence of the ring index (RI) for all tetraethers (GDGT, GTGT, and GMGT), GDGT, and GMGT (RI ± standard deviation). (G) Strain dependence of the GMGT/GDGT ratio.

## Discussion

### The membrane response of *Pyrococcus furiosus* follows a particular strategy involving the alteration of the GMGT/GDGT ratio

The 14 different core lipid structures identified in the present study in the hyperthermophilic and neutrophilic marine archaeon *Pyrococcus furiosus* are in accordance with those reported previously (Tourte *et al.*, 2020a), which included DGD, GTGT0 to 2, GDGT0 to 4 and GMGT0 to 4 (Figure S1). GTGT2 and GMGT2 to 4 were only sporadically reported in Archaea (see for instance Knappy *et al.*, 2011; Bauersachs *et al.*, 2015 and references therein), but were regularly detected in *P. furiosus,* confirming the very peculiar core lipid composition of this archaeon compared to closely related Thermococcales and to other Archaea (Tourte *et al.*, 2020b). Here, we examined the role of these peculiar lipids in the stress response of *P. furiosus,* and showed that all but the presence/absence of sulfur in the growth medium affected the core lipid composition of *P. furiosus* to some extent.

In contrast to other archaea producing both di- and tetraether lipids (Lai *et al.*, 2008; Matsuno *et al.*, 2009; Cario *et al.*, 2015), we have surprisingly found no evidence for a regulation of the diether/tetraether ratio in *P. furiosus.* In addition, although a trend seems to exist between the RI and some of the stressors tested, the variations in the number of rings per molecule remain very limited (often below 0.2), and do not reach what is observed for instance in thermoacidophiles, which RI values usually vary by *ca.* 0.5-1 unit over the range of the environmental stressor (Shimada *et al.*, 2008; Feyhl-Buska *et al.*, 2016). *P. furiosus* thus appears to rely on an uncommon and specific membrane adaptation strategy which involves the modification of the ratio of two tetraether core lipid classes, namely glycerol mono- and dialkyl glycerol tetraethers (GMGT and GDGT, respectively).

GMGT are a particular type of tetraether lipids exhibiting a covalent C-C bond between the two C40 alkyl chains (Morii *et al.*, 1998) and can represent a significant proportion of membrane lipids in numerous archaea, such as *Methanothermus fervidus* (31 %; Morii *et al.*, 1998)*, Ignisphaera aggregans* (39 %; Knappy *et al.*, 2011), *Methanothermococcus okinawensis* (15 %; Baumann *et al.*, 2018), and a few Thermococcales (e.g., 50 % in *Thermococcus waiotapuensis,* 34 % in *P. horikoshii,* and *ca.* 15 % in *T. celer* and *T. guaymasensis;* Sugai *et al.*, 2004; Tourte *et al.*, 2020b). Since all these archaea are (hyper)thermophiles and since the abundance of GMGT was observed to positively correlate with the mean annual air temperature in peats (Naafs *et al.*, 2018), GMGT were first associated with the adaptation to increased temperature. Although the role of GMGT in terms of membrane structure and properties remains uncharacterized, the presence of a covalent C-C bond between the two alkyl chains suggests that the free motion of the GMGT molecule will be strongly reduced, thus resulting in increased membrane stability and impermeability compared to GDGT in a manner similar to that of cMGD *vs.* DGD (Sprott *et al.*, 1991; Dannenmuller *et al.*, 2000; Arakawa *et al.*, 2001). Building on this hypothesis, GMGT were then proposed to partake in adaptation of Archaea to high temperature, but also to high salinity and low pH (Schouten *et al.*, 2008a; Knappy *et al.*, 2011). However, to date, there has been no demonstration of the role of GMGT in the membrane response of Archaea to any stressor, which thus remains elusive. While GMGT did appear to have a role in adaptation to temperature in *P. furiosus*, their proportion relative to GDGT decrease when the strain was grown at high temperatures, e.g., 43.4 ± 3.1 % at 85 °C *vs.* 20.8 ± 2.0 % at 98 °C (Figure 2G, Table 2), which is opposite to the trend previously reported for these compounds. Such variations are also antagonistic with the proportion of cMGD relative to DGD in *M. jannaschii*, which rose with increasing temperature, e.g., 12 % at 44 °C to *ca.* 45 % at 65 °C (Sprott *et al.*, 1991). The relative amounts of bilayer-forming cMGD in *M. jannaschii* and monolayer-forming GMGT in *P. furiosus* being opposite in response to growth temperature clearly indicate that the membranes behave in fundamentally distinct ways under temperature stress. Further characterization of their physicochemical properties is now sorely required to elucidate the features of the membranes they create and to rationalize their adaptive functions in response to temperature.

In contrast to temperature, relatively little is known about the adaptive strategies Archaea implement in response to pH and salinity stress outside of thermoacidophiles and extreme halophiles, respectively. One would nonetheless expect the membrane impermeability to ions and water to increase with osmotic or proton pressures in order to ensure proper cellular functioning and integrity. No cMGD has been detected so far in halophilic archaea, which instead shield their membrane with negatively charged polar head groups in order to preserve membrane impermeability under extreme salt conditions (Tenchov *et al.*, 2006; Kellermann *et al.*, 2016). Our procedure for core lipid extraction resulting in the excision of the polar head groups, such a strategy could not be investigated here. The adaptation to pH of thermoacidophilic archaea to pH implementing a modulation of the number of cyclopentane rings in tetraether lipids (Shimada *et al.*, 2008; Feyhl-Buska *et al.*, 2016) could on the other hand be assessed here. No significant modification of the number of cyclopentane rings was observed in the tetraether pool of *P. furiosus* with changing pH or salinity (Table 1, Figure 3F and Figure 4F). However, as aforementioned, the covalent C-C bond between the two alkyl chains in GMGT is supposed to provide a more efficient barrier to solutes and water than the classic DGD and GDGT, respectively. One would thus expect an increase of the GMGT/GDGT ratio with increasing salinity and decreasing pH, which was exactly what was observed for *P. furiosus* (Figure 3G and Figure 4G). These results suggest that, at least for *P. furiosus,* GMGT are essential lipids for the membrane adaptation to salinity and pH, and could for instance help maintaining proper membrane permeability to solutes and protons under stressful conditions. Such an adaptive function again contrasts with environmental data, for which a positive correlation between pH and the relative abundance of GMGT has been observed in both low temperature (peats, < 20 °C; Naafs *et al.*, 2018) and high temperature (terrestrial hydrothermal vents, > 50 °C; Jia *et al.*, 2014) settings. Elucidating the adaptive functions of GMGT in response to pH and salinity in other archaeal species, such as *Ignisphaera aggregans*, a freshwater neutrophilic hyperthermophile (Knappy *et al.*, 2011), or *Aciduliprofundum boonei,* a marine acidophilic hyperthermophile (Schouten *et al.*, 2008), is now essential to determine whether the mechanisms observed in *P. furiosus* are specific to this marine neutrophilic hyperthermophile or shared between ecologically and phylogenetically distant archaea.

### The regulation of the number of cyclopentane rings in membrane lipids is a minor component of the adaptive response in *Pyrococcus furiosus*

Tetraether lipids form monolayer membranes that are more rigid and impermeable than typical bilayer membranes (Chong, 2010). The presence of one to eight cyclopentane rings further enhances the packing of the monolayer (Gliozzi *et al.*, 1983; Gabriel and Chong, 2000; Chong, 2010), thus reducing proton and solute permeability and maintaining membrane stability at high temperatures. Ring-containing GDGT were initially identified in thermoacidophilic archaea, such as *Sulfolobus acidocaldarius* (De Rosa *et al.*, 1983) or *Thermoplasma acidophilum* (Shimada *et al.*, 2002), although these compounds were more recently demonstrasted to be also vastly distributed within mesophilic archaea, such as Thaumarchaeota (e.g., Schouten *et al.*, 2008b; Elling *et al.*, 2017). Several studies have shown that the membrane adaptive response to temperature triggers an increase of the RIGDGT in thermoacidophiles, such as *Acidilobus sulfurireduscens* (+0.6 cycle from 65 °C to 81 °C; Boyd *et al.*, 2011), but also in mesophiles, such as *Nitrosopumilus maritimus* (+0.7 cycle from 22 °C to 28 °C; Elling *et al.*, 2015), suggesting that the regulation of the RIGDGT in Archaea is a common membrane adaptation strategy to increased temperature (Uda *et al.*, 2004; Feyhl-Buska *et al.*, 2016; Bale *et al.*, 2019). Modulations of the RI_GDGT_ have also been observed in thermoacidophiles grown at varying pH (Boyd *et al.*, 2013; Feyhl-Buska *et al.*, 2016). However, whether and how these variations contribute to the membrane response to pH remains unclear as the RIGDGT decreases with pH for some species (−0.6 cycles from pH 3.0 to pH 5.0 in the case of *Acidilobus sulfurireduscens;* Boyd *et al.*, 2011) while it increases with pH for some others (e.g., +1.1 cycles from pH 1.2 to pH 3.0 for *Thermoplasma acidophilum;* Shimada *et al.*, 2008), thus challenging the predictions based on the physicochemical properties of these lipids. To our knowledge, only one study investigated the variations of the RI_GDGT_ in response to varying salinity and reported no significant modification with the environmental stressor (Elling *et al.*, 2015).

In *P. furiosus,* RI values were also significantly positively correlated with temperature (Table 2 and Figure 2F). The temperature of 103 °C was notably the only one at which we could detect GDGT4 (Figure 2B). Interestingly, the RI_GMGT_ followed the same trend (Figure 2F), suggesting that the regulation of the RI in response to temperature is independent of the tetraether core structure. In contrast to mesophilic neutrophiles which do not modulate their RI_GDGT_ in response to pH (Elling *et al.*, 2015), *P. furiosus* significantly increased the RI_GDGT_ with increasing pH (Figure 3F, Tables 1 and 2). In contrast, both RI_GDGT_ and RI_GMGT_ varied significantly only at the highest salinity tested (Figure 4F), suggesting that the decrease of the RI in response to increasing salinity could be triggered only above a salinity threshold. Despite these observations, the low RI in *P. furiosus* (< 0.6 under all the conditions tested) as compared to that of other archaea (generally > 1) indicates that the regulation of the average number of cyclopentane rings in tetraether lipids may be negligible in the adaptive strategy of *P. furiosus.*

### Diether lipids are not involved in membrane adaptation in *Pyrococcus furiosus*

Diether lipids form bilayer membranes that are more fluid and permeable than their tetraether-based monolayer counterparts (Chong, 2010). They thus play a major role in the membrane adaptation in Archaea capable of synthesizing both diether and tetraether lipids (Sprott *et al.*, 1991; Matsuno *et al.*, 2009; Cario *et al.*, 2015). While some variations of the DGD relative abundances did occur in *P. furiosus* (Table 1), no significant differences nor correlations with temperature or any other parameter tested were observed in the present study (Table 2), suggesting that, in contrast to other Thermococcales, the modulation of the diether/tetraether ratio might not be part of the adaptive response of *P. furiosus*. This however does not necessarily preclude the existence of adaptive functions for diether lipids in *P. furiosus.* Indeed, we previously reported that in Thermococcales, diether lipids can harbor up to seven different polar head groups of variable size and polarity in contrast to tetraether lipids which all harbor two phosphoinositol headgroups (Tourte, *et al.*, 2020a). Although no adaptive response could be detected for diether core lipids in the case of *P. furiosus*, the possibility that diether lipids could respond through the alteration of their polar head groups cannot be ruled out. Regardless, the presence of relatively large proportions of diether lipids in *P. furiosus* is puzzling since the stability of diether lipid-based membranes at its optimal growth temperature of 98 °C remains questionable, and suggests that diether lipids could harbor other important physiological functions. They may for instance only exist in mixture with tetraether lipids, i.e., diether lipids would be dispersed throughout a monolayer membrane, and, given their very divergent properties, act as membrane fluidizing agents in *P. furiosus.* However, no data about the spatial distribution of both types of lipids in the archaeal membrane have been reported to date, and these functions of diether lipids in the membrane of *P. furiosus* thus remain hypothetical.

### The carbon source and the genetic content strongly affect the lipid composition of *Pyrococcus furiosus*

It is now well documented that Archaea regulate their lipid composition in response to various parameters besides temperature, pH, and salinity, e.g., carbon, phosphorous, and nitrogen sources and availability, growth rate, and oxygen content (Langworthy, 1977; Matsuno *et al.*, 2009; Elling *et al.*, 2014; Meador *et al.*, 2014; Feyhl-Buska *et al.*, 2016; Quehenberger *et al.*, 2020; Zhou *et al.*, 2020). Here, we also tested the impact of the carbon source by comparing growth in DC (disaccharides) and TRM (polypeptides) media. Growth in the DC medium boosted the synthesis of ring-containing tetraethers to such an extent that *P. furiosus* exhibited RI values similar to those observed for Thaumarchaeota and thermoacidophilic archaea (Figure 5F). Interestingly, this mimics the response of *S. acidocaldarius* grown under nutrient limitation (Bischof *et al.*, 2019), suggesting that the RI increase observed in *P. furiosus* grown in DC medium could similarly result from energy flux slowdown. Unfortunately, no single monosaccharide can support growth of *P. furiosus* and only disaccharides could be used as carbohydrate carbon source in DC medium, preventing the identification of the limiting reaction which might for instance be the cleavage of di- into monosaccharides or the breakdown of monosaccharides. The shift in RI observed in DC medium appeared much larger than those observed with salinity, pH, and temperature variations (Table 2), indicating that metabolism rate limitation, which is very frequent in the natural environment, may be a greater environmental stressor for *P. furiosus* than any other parameter tested here, an observation congruent with that made in other archaea (Hurley *et al.*, 2016).

Additionally, we compared the lipid composition of two quasi-isogenic strains, i.e., the wild-type strain DSM3638 and its derivative strain COM1. Although strain COM1 differs from strain DSM3638 by limited genomic modifications (Bridger *et al.*, 2012), it exhibited a completely distinct core lipid composition. For instance, COM1 and DSM3638 showed significantly different GDGT and GMGT proportions (Figure 6A-C). In addition, as seen for the wild-type strain in DC medium, strain COM1 showed much higher RI than strain DSM3638 (Figure 6F). Modification of the genomic region near the gene responsible for cyclization could directly explain the increase of the RI of strain COM1. However, the genomic comparison of the two strains did not allow the identification of any noticeable change in the vicinity of gene PF0210, which is homologous to the recently identified two radical S-adenosylmethionine (SAM) proteins involved in GDGT cyclization (Grs) in *Sulfolobus acidocaldarius* (Zeng *et al.*, 2019) and might similarly be involved in the formation of the GMGT interchain C-C bond. These differences in RI might also result from uracil starvation as seen in *Sulfolobus acidocaldarius* (Bischof *et al.*, 2019), since strain COM1 is an uracil auxotroph while strain DSM3638 is an autotroph (Lipscomb *et al.*, 2011). Overall, it is highly probable that these notable differences in lipid composition result from the general disturbance of the cell regulation network triggered by the few chromosomal rearrangements present in COM1. Chromosome stability in Thermococcales and other Archaea is a highly debated issue. It is influenced by numerous intrinsic factors, e.g., the genomes of *Pyrococcus* species are highly rearranged (Zivanovic, 2002; Cossu *et al.*, 2015), or extrinsic factors, such as virus infection or mobile element insertion (Cossu *et al.*, 2017), that could strongly affect the lipid composition of their host. Altogether, our results indicate that besides typical abiotic factors, numerous biotic factors also greatly impact the average number of cyclopentane rings and the overall lipid composition in Archaea.

## Material and methods

### Microorganism and growth conditions

*Pyrococcus furiosus* strain DSM3638 was purchased from the Deutsche Sammlung von Mikroorganismen und Zellkulturen (DSMZ, Braunschweig, Germany). *Pyrococcus furiosus* strain COM1 was kindly provided by the Adams lab (University of Georgia, Athens, Georgia, USA). It is a naturally competent derivative of DSM3638 that has been developed for genetic manipulations by deleting its *pyrF* gene to make it auxotroph for uracil (Lipscomb *et al.*, 2011; Farkas *et al.*, 2012). Strain COM1 exhibits several chromosomal rearrangements, deletions, insertions, and single base modifications compared to the type strain DSM3638 (Bridger *et al.*, 2012).

Cultures were routinely grown under optimal growth conditions, i.e., at 98 °C and pH 6.8, with 3 % *w/v* NaCl and 10 g. L^-1^ elemental sulfur, in a rich medium established for Thermococcales (TRM; Zeng *et al.*, 2009), containing for 1 L: MgCl_2_, 5 g; peptone (Difco), 4 g; PIPES [piperazine-N,N’-bis(2-ethanesulfonic acid)], 3.3 g (10 mM); yeast extract (Difco), 1 g; KCl 0.7 g; (NH_4_)_2_SO_4_ 0.5 g; KH_2_PO_4_ 50 mg; K_2_HPO_4_ 50 mg; NaBr 50 mg; CaCl_2_ 20 mg; SrCl_2_ 10 mg; FeCl_3_ 4 mg; Na_2_WO_4_ 3 mg and resazurin (Sigma Aldrich) 1 mg. Alternatively, cultures were grown in DC medium (Lipscomb *et al.*, 2011), a defined medium with cellobiose as a carbon source at 2.8 % *w/v* NaCl and pH 6.8, containing for 1 L: MgSO_4_, 3.5 g; cellobiose (Alfa Aesar), 3.5 g; MgCl_2_, 2.7 g; cysteine-HCl (Sigma Aldrich), 1 g; NaHCO_3_, 1 g; KCl, 0.3 g; NH_4_Cl, 250 mg; CaCl_2_, 140 mg; KH_2_PO_4_, 140 mg; K_2_HPO_4_, 170 mg; Na_2_WO_4_, 0.3 mg, amino acid solution, 40 mL; vitamin solution, 5 mL; and trace mineral solution, 1 mL. Growth media were supplemented with uracil (20 μM final concentration) for *P. furiosus* strain COM1. Strict anaerobiosis was ensured by addition of Na_2_S (0.1 % *w/v* final concentration) before inoculation.

We evaluated the membrane response of *P*. *furiosus* to the following parameters: temperature (80, 85, 90, 98, and 103 °C), salinity (1, 2, 3, and 4 % *w/v* NaCl), presence/absence of elemental sulfur (0 and 10 g L^-1^), type of growth medium (TRM and DC) and genetic background (the wild-type strain DSM3638 *vs*. strain COM1). Growth of *P. furiosus* was first reported at pH values ranging from 5 to 9 using only 0.05 M of glycylglycine as buffer for pH ≥ 8.0 (Fiala and Stetter, 1986). In contrast, we could not maintain pH ≥ 6.8 by adding up to 1 M of either glycylglycine, 2-amino-2-methyl-1-propanol (AMP), 2-Amino-2-methyl-1,3-propanediol (AMPD), 3-(cyclohexylamino)-2-hydroxy-1-propanesulfonic acid (CAPSO), 2- (cyclohexylamino)ethanesulfonic acid (CHES), or PIPES buffer, likely because of the production and excretion of organic acids during growth. Monitoring growth in alkaline cultures showed that growth essentially picked up only after pH was lowered to values close to the optimum. Thus, only endpoint pH values are reported here (5.5, 5.6, 5.9, 6.2, 6.4 and 6.6). Cultures were set up with 10 mM PIPES for pH ≥ 6.0, and 10 mM 2-(N-morpholino)ethanesulfonic acid (MES) for pH ≤ 6.0. Growth was monitored by counting with a Thoma cell counting chamber (depth 0.01 mm) under a light microscope (Thermo Fisher Scientific EVOS® XL Core 400×, Waltham, MA, USA), and growth curves were established under each condition on at least three biological replicates (for evaluation of the growth under each condition, refer to Table S1).

### Core lipid extraction and HPLC-MS analysis

Cells of 250-mL cultures in late exponential phase were recovered by centrifugation (4000 × g, 45 min, 4 °C) and rinsed twice with an isotonic saline solution. The cell pellets were lyophilized overnight and kept at −80 °C until lipid extraction. Extraction was performed on three biological replicates. Following acid hydrolysis of the dried cells (1.2 N HCl in methanol at 110 °C for 3 h), core lipids were extracted by filtration over celite using methanol/dichloromethane (1:1, *v*/*v*). The resulting solvent extracts were dried under reduced pressure, solubilized in *n*-heptane/isopropanol (99:1, *v*/*v*) and analyzed by high-performance liquid chromatography-mass spectrometry (HPLC-MS) using an HP 1100 series HPLC instrument equipped with an auto-injector and a Chemstation chromatography manager software connected to a Bruker Esquire 3000^Plus^ ion trap mass spectrometer, as described in (Tourte *et al.*, 2021). A standard solution containing core DGD and GDGT0 in a 2/1 molar ratio demonstrated a molar response factor of DGD *ca*. 10 times lower than that of GDGT0 under our analytical conditions. In the absence of a measured response factor for the different tetraether lipids, we assumed that all tetraethers have the same response factor as GDGT0. Core lipid relative abundances were determined by integration of the peak area on the mass chromatograms corresponding to the [M+H]^+^ ion of the different core lipids using a Bruker Data Analysis mass spectrometry software (version 4.2), and the relative abundances of DGD relative to that of tetraethers were corrected by a factor of 10.

### Statistical analyses

Statistical analyses were computed using the functions implemented within the base R core package (version 3.6.3; R Core Team, 2020). The weighted average number of rings per lipid molecule, or ring index (RI), was calculated as follows (Schouten *et al.*, 2007):

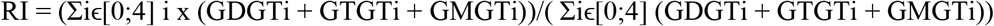

Data normality and homoscedasticity were assessed using the Shapiro-Wilk and Levene tests, respectively. Lipid relative abundances under each condition were compared using the Student t-test when there were only two independent groups of conditions, i.e., presence of sulfur, medium and strain. With more than two groups, comparisons were computed using one-way ANOVA and Tukey tests (normality and homoscedasticity of the data; % NaCl) or Kruskal-Wallis and Dunn tests (data not normally distributed and significantly different variances; temperature and pH). Correlations between the lipid proportions and temperature, pH, and salinity were assessed by the two-tailed probability associated with the Spearman correlation coefficient (ρ). Differences and correlations with *P*-values below 0.05 were considered significant.

## Acknowledgments

MT is supported by a Ph.D. grant from the French Ministry of Research and Technology. The authors would like to thank the French National Research Agency for funding the ArchaeoMembranes project to PO (ANR-17-CE11-0012-01) and the CNRS Interdisciplinary program ‘Origines’ for funding the ReseArch project. The authors gratefully acknowledge Prof. Michael W.W. Adams for kindly providing *P. furiosus* strain COM1.

