## Supplementary material for "Membrane adaptation in the hyperthermophilic archaeon *Pyrococcus furiosus* relies upon a novel strategy involving glycerol monoalkyl glycerol tetraether lipids"

#### Supplementary figures

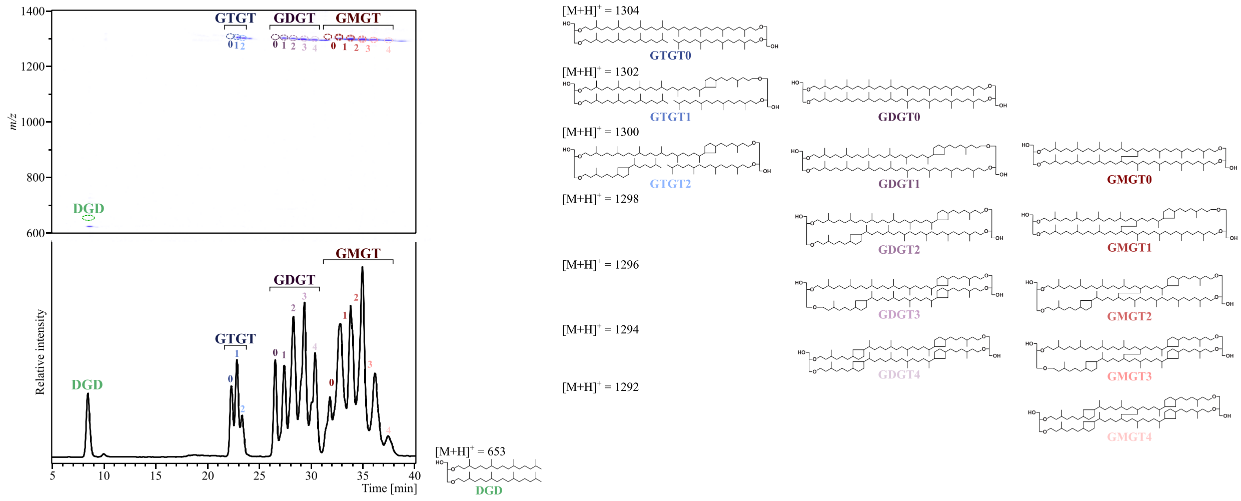


**Figure S1: *Pyrococcus furiosus* synthesizes a large diversity of tetraether core structures.**
Core lipids were detected in positive ion mode. The density map and chromatogram from *P. furiosus* grown in DC medium are displayed (zoom in the 5-40 min, *m/z* 600-1400 window). The HPLC chromatogram was drawn by extracting the following protonated ion masses with a mass deviation of ± 0.5 Da: 653.7, 1294.2, 1296.2, 1298.2, 1300.2, 1302.2, 1304.2. Note that two isomers were detected for all GMGTs, except GMGT4, resulting in doubled, partially overlapping peaks. Shorthand nomenclature and protonated mass-charge ratios are indicated for each structure. Core structures: dialkyl glycerol diethers (DGD; green), glycerol dialkyl glycerol tetraethers with 0 to 4 cyclopentane rings (GDGT0 to 4; dark to light purple), glycerol trialkyl glycerol tetraethers with 0 to 2 cyclopentane rings (GTGT0 to 2; dark to light blue) and glycerol monoalkyl glycerol tetraethers with 0 to 4 cyclopentane rings (GMGT0 to 4; dark to light red). The same color code is used throughout the whole manuscript. Position of the cyclopentane rings and of the covalent C-C bond between the two alkyl chains are drawn arbitrarily.

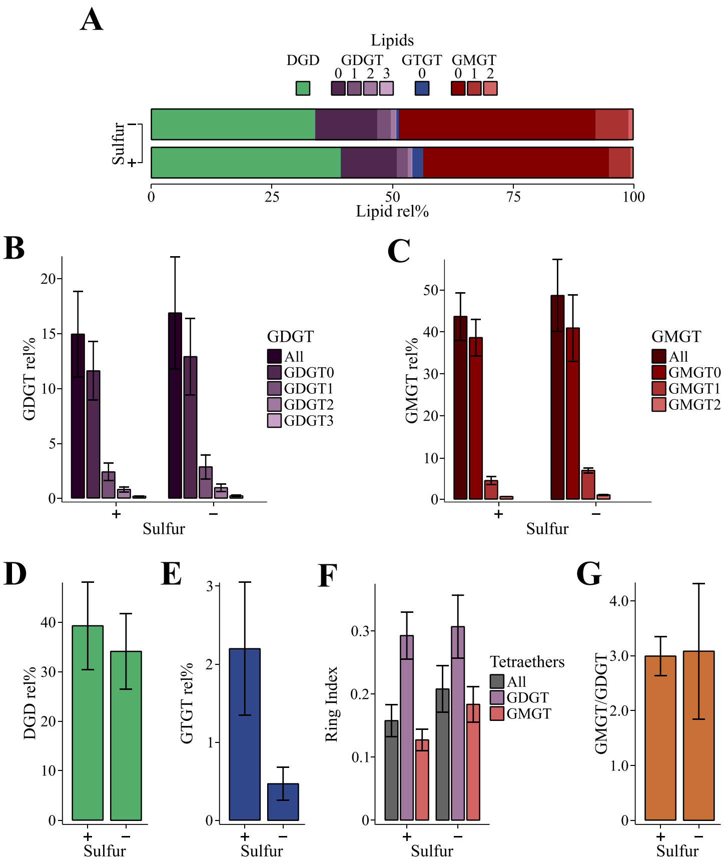


**Figure S2: The presence of elemental sulfur does not trigger modifications of the membrane lipid composition in *Pyrococcus furiosus*.**

*P. furiosus* DSM3638 was grown under optimal conditions (TRM at 98 °C, pH 6.8, 3% *w/v* NaCl) with 0 (-S) and 10 g $L^{-1}$ (+S) elemental sulfur. +S represents the optimal growth condition. Only the core structures identified are indicated. Error bars represent the standard deviation calculated on three biological replicates. (A) Total core lipid compositions under each sulfur condition. (B) Influence of the presence of sulfur on GDGT relative proportions. GDGT (dark purple) corresponds to the summed GDGT, regardless of the cyclopentane ring content. (C) Influence of the presence of sulfur on GMGT relative proportions. GMGT (dark red) corresponds to the summed GMGT, regardless of the cyclopentane ring content. (D) Influence of the presence of sulfur on DGD relative abundance. (E) Influence of the presence of sulfur on GTGT0 relative abundance. (F) Elemental sulfur dependence of the ring index (RI) for all tetraethers (GDGT, GTGT, and GMGT), GDGT, and GMGT (RI ± standard deviation). (G) Elemental sulfur dependence of the GMGT/GDGT ratio.

### Supplementary tables

**Table S1.** ***Pyrococcus furiosus*** **growth parameters under each condition tested here (averaged over 3 biological replicates ± standard deviation).**

| | | **Doubling time (h)** | **Harvest time (h)** | **Harvest cell count (× $10^8$; cell $ml^{-1}$)** |
|---|---|---|---|---|
| Temp. (°C) | 80 | 6.83 ± 0.33 | 69.32 ± 3.33 | 1.13 ± 0.23 |
| | 85 | 3.23 ± 0.54 | 33.62 ± 5.60 | 1.37 ± 0.12 |
| | 90 | 2.55 ± 0.17 | 28.41 ± 1.86 | 2.25 ± 0.71 |
| | 98* | 2.50 ± 0.21 | 29.26 ± 2.41 | 2.93 ± 1.29 |
| | 103 | 1.75 ± 0.08 | 18.48 ± 0.88 | 1.55 ± 0.64 |
| pH | 5.5 | 5.63 ± 0.70 | 56.84 ± 3.75 | 1.10 ± 0.10 |
| | 5.6 | 5.35 ± 0.24 | 54.41 ± 2.44 | 1.15 ± 0.93 |
| | 5.9 | 3.85 ± 0.96 | 41.64 ± 10.41 | 1.80 ± 0.36 |
| | 6.2 | 2.67 ± 0.30 | 28.45 ± 3.18 | 1.60 ± 0.36 |
| | 6.4 | 2.83 ± 0.72 | 31.08 ± 7.95 | 2.03 ± 0.81 |
| | 6.6* | 2.50 ± 0.21 | 29.26 ± 2.41 | 2.93 ± 1.29 |
| %NaCl (%w/v) | 1 | 2.97 ± 0.31 | 32.15 ± 3.39 | 1.83 ± 0.76 |
| | 2 | 2.68 ± 0.37 | 29.90 ± 4.01 | 2.27 ± 0.68 |
| | 3* | 2.50 ± 0.21 | 29.26 ± 2.41 | 2.93 ± 1.29 |
| | 4 | 3.13 ± 0.35 | 31.86 ± 3.62 | 1.17 ± 0.29 |
| Sulfur | +S* | 2.50 ± 0.21 | 29.26 ± 2.41 | 2.93 ± 1.29 |
| | -S | 3.65 ± 0.23 | 37.97 ± 2.38 | 1.47 ± 0.35 |
| Medium | TRM* | 2.50 ± 0.21 | 29.26 ± 2.41 | 2.93 ± 1.29 |
| | DC | 3.33 ± 0.22 | 35.21 ± 2.30 | 1.53 ± 0.25 |
| Strain | DSM3638* | 2.50 ± 0.21 | 29.26 ± 2.41 | 2.93 ± 1.29 |
| | COM1 | 2.82 ± 0.17 | 29.02 ± 1.80 | 1.25 ± 0.62 |

*Optimal growth conditions used as reference.
